# Advancing Plant Metabolic Research By Using Large Language Models To Expand Databases And Extract Labelled Data

**DOI:** 10.1101/2024.11.05.622126

**Authors:** Rachel Knapp, Braidon Johnson, Lucas Busta

## Abstract

Premise: Recently, plant science has seen transformative advances in scalable data collection for sequence and chemical data. These large datasets, combined with machine learning, revealed that conducting plant metabolic research on large scales yields remarkable insights. A key next step in increasing scale has been revealed with the advent of accessible large language models, which, even in their early stages, can distill structured data from literature. This brings us closer to creating specialized databases that consolidate virtually all published knowledge on a topic. Methods: Here, we first test different prompt engineering technique / language model combinations in the identification of validated enzyme-product pairs. Next, we evaluate automated prompt engineering and retrieval augmented generation applied to identifying compound-species associations. Finally, we build and determine the accuracy of a multimodal language model-based pipeline that transcribes images of tables into machine-readable formats. Results: When tuned for each specific task, these methods perform with high accuracies (80-90 percent for enzyme-product pair identification and table image transcription), or with modest accuracies (50 percent) but lower false-negative rates than previous methods (down to 40 percent from 55 percent) for compound-species pair identification. Discussion: We enumerate several suggestions for working with language models as researchers, among which is the importance of the user’s domain-specific expertise and knowledge.

**Significance Statement:** Scientific databases have played a major role in advancing metabolic research. However, even today’s advanced databases are incomplete and/or are not built to best suit certain research tasks. Here, we explored and evaluated the use of large language models and various prompt engineering techniques to expand and subset existing databases in task-specific ways. Our results illustrate the potential for high-accuracy additions and restructurings of existing databases using language models, assuming the specific methods by which the models are used are tuned and validated for the specific task. These findings are important because they outline a method by which we could greatly expand existing databases and rapidly tailor them to specific research efforts, leading to greater research productivity and effective utilization of past research findings.

All authors collected data, analyzed data, prepared the manuscript, and approved its final version. The authors declare that they have no competing interests.

## 1. Introduction

Public databases are powerful tools that researchers can use to enhance studies of plant chemical diversity and metabolism. Significant efforts have been made to develop and populate resources such as Phytozome, GenBank, and UniProt, which support research into plant genomes and proteomes (1–3); The Plant Metabolic Network and KEGG, which support metabolic research (4, 5); GNPS, MassBank, and MassBank of North America, which support mass spectrometry research (6, 7); and Natural Products Online’s LOTUS, which aids in studying the distribution of plant natural products across phylogenies (8). These databases are essential for foundational tasks in plant metabolic research, such as identifying biosynthetic enzymes and tracking their evolution, identifying compounds in plant extracts, and understanding how plant products are distributed across the plant kingdom. Despite their utility, however, these databases remain incomplete. Much data remains locked in PDFs of scientific articles or in supplementary files, and current databases may not meet users’ specific needs for data structure or curation, particularly in lineages that lack well-studied model or crop plants. This contributes to challenges researchers face when assembling, refining, and curating the information necessary for project-specific analyses. Addressing these limitations is crucial, otherwise researchers will continue to face ongoing obstacles in leveraging existing data, likely hindering advances in plant biochemistry, natural product discovery, and metabolic engineering.

To overcome challenges related to data mining and database structures, researchers from diverse fields are increasingly using large language models. These models serve a variety of purposes: embedding models transform sequences (such as words or proteins) into numerical vectors for analysis; chat/completion models generate coherent, context-aware text; classification models label input data based on predefined categories; and multimodal models integrate multiple data types, like text and images, for tasks such as image captioning. Some models can also follow user-defined prompts - text-based instructions that provide detail on how the language model should perform a given task. Many recent studies report on prompt engineering strategies to improve the accuracy of a model’s output. Examples include few-shot and multi-shot learning, where the model is given one or more examples of question-answer pairs, and techniques like chain-of-thought and Tree of Thought, which guide the model through step-by-step reasoning for complex tasks (9, 10). Inspired by these approaches, researchers in the natural sciences, including the field of plant chemistry, are already using language models to tackle data mining and database-related problems. For instance, some are summarizing large literature landscapes (11), while others are extracting information about enzyme-substrate pairs (12), linking specific phytochemicals to specific plant species through data mining (13), and identifying yet unreported enzyme activities from relatively understudied domains of life (14). These examples illustrate the transformative potential of language models and their ability to advance our field, but also highlight that we are at the very beginning when it comes to integrating language models into plant metabolic research.

Here, we provide several examples of how language models can help plant scientists address challenges related to extracting and organizing data from unstructured or loosely structured sources. We also report results from experiments with language models of different levels of sophistication (including models that can process images) as well as with various prompt engineering strategies. First, we used prompt engineering to extract enzyme-product relationships from NCBI entries, enabling the separation of validated enzyme-product pairs from predicted enzyme-product pairs. Next, we developed an automated prompt engineering strategy to improve the extraction of chemical compound-plant species relationships from text. Finally, we integrated vision models with prompts and language model calls to extract structured data from PNG images of tables in research article PDF files. Our approaches achieved high levels of accuracy and illustrate the potential of language models in data organization projects. The present work also highlights the critical importance of model users’ detailed knowledge and subject matter expertise in plant metabolism.

## 2. Materials and Methods

Enzyme and compound information used as data in Section 3A was retrieved from NCBI via the NCBI API and processed in R using the rentrez package (15). For data handling and visualization, tidyverse (16) and ggplot2 (17) were used together with ggupset (18) for plotting. All significance testing was conducted using the statix package (19). The completion language models used in this study were OpenAI models accessed via the OpenAI API. In Section 3A we manually developed system prompts for these models based on iterative testing of the prompts’ performance. For automated prompt engineering in Section 3B, we used gpt-4 to generate prompts, using WandB (20) to automate their creation and test prompt performance across a manually curated set of previously published test cases (13). Embeddings were created with the BAAI/bge-small-en-v1.5 embedding model downloaded from HuggingFace (https://huggingface.co/BAAI/bge-small-en-v1.5). Retrieval of passages for the RAG system were obtain by having gpt-4 generate a few sentences resembling an academic answer to the question, converting those into embeddings, then using a nearest-neighbors algorithm (sklearn.neighbors.NearestNeighbors) to find the most similar matches in the vector database of abstracts. For the Vision AI tasks in Section 3C, several Python modules were used to help transcribe and process table images by turning them into structured JSON formats including os for file handling, base64 for image encoding, requests for API interactions, pandas for data manipulation, and openai for model integration. The PNG images of tables were base64-encoded and analyzed using OpenAI’s gpt-4o model. That same model was used to check if both plant species and compound names were present in the intial transcriptions and to determine the table’s orientation—whether rows represented species or compounds—to ensure accurate data extraction. The R package ggtree was used to visualize the phylogenetic tree (21). All code and raw data used in this project are available at https://github.com/thebustalab/ai_in_phytochemistry.

## 3. Results and Discussion

### A. Extracting Validated Enzyme-Product Relationships

The goal in this subsection was to assess the ability of large language models to distinguish between database records that report the validated product of an enzyme catalyst (positive entries) versus those that report a predicted product (negative entries). The first step towards this goal was to obtain a collection of database records with both positive and negative representatives. To obtain this collection we first conducted command-line NCBI searches using three enzyme-related search terms (“beta-amyrin synthase”, “lupeol synthase”, “cycloartenol synthase”) and added the first 20 records from each search into our collection, resulting in an initial set of 60 records identifed by their protein IDs (Fig. 1A). Manual inspection of these 60 records revealed an imbalance, with a predominance of negative records (reports of enzymes and their predicted, not validated, products). To create a balanced collection, we supplemented the initial set of 60 with additional records that we identified based on manual inspection of peer-reviewed articles that reported the validated products of specific enzymes. After these steps our collection consisted of 142 records, including 93 positive records and 49 negative records.

**Fig. 1.**
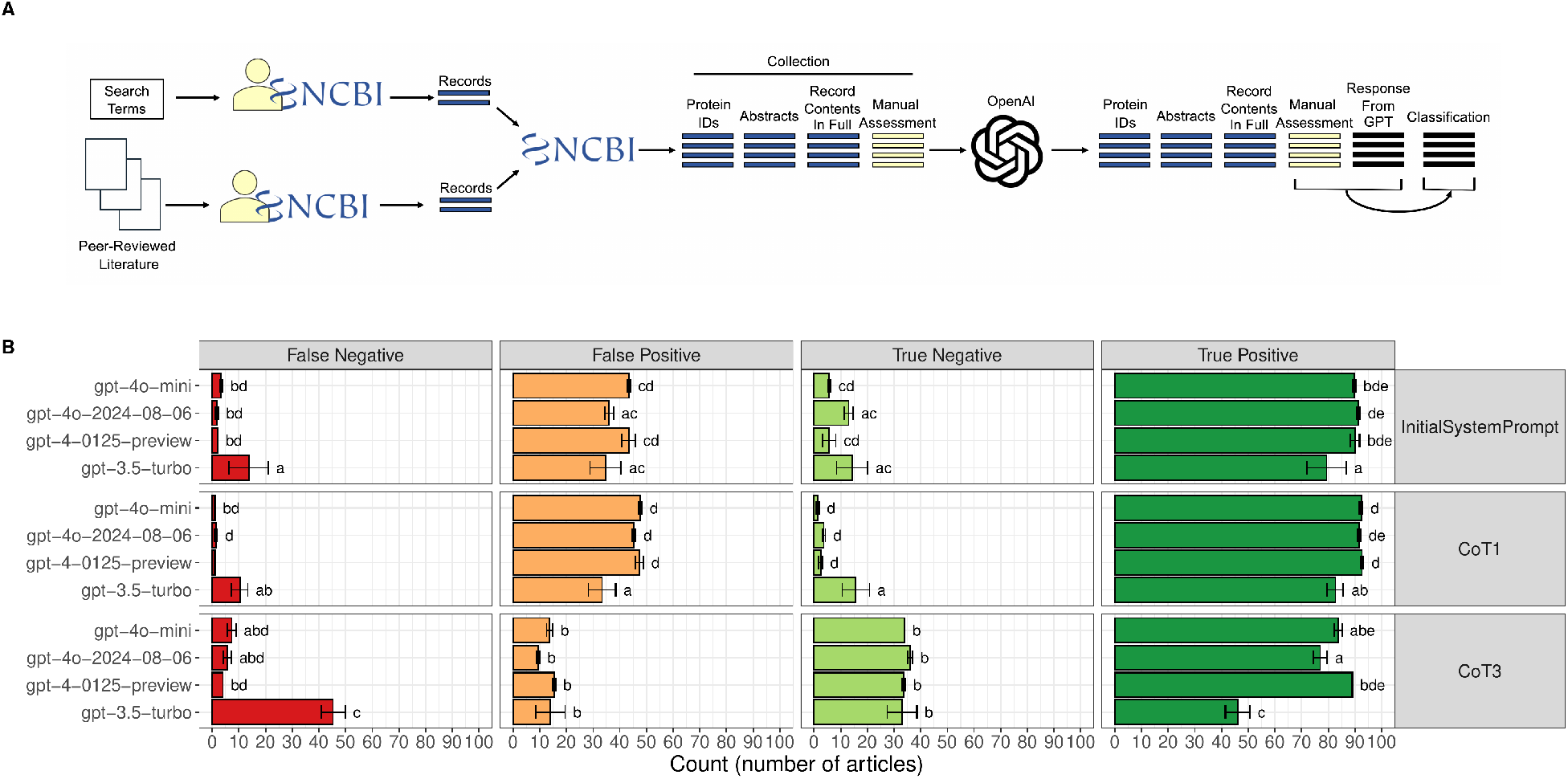
Workflow and results of protein-product relationship evaluation. **A** Workflow used to generate results. Search terms indicate enzymes searched for in NCBI (for example, “beta-amyrin synthase” or “cycloartenol synthase”). Peer-reviewed literature indicates the manually curated database of NCBI protein entries that describe verified protein-product pairs. The collection is made up of both the products of the search terms and the contents of the curated database. Blue lines indicate NCBI protein records, yellow lines indicate cells created through manual evaluation, black boxes represent cells generated via language model response and comparison between language model response and manual evaluation. **B** Outcomes of language model classification of protein-product pairs. The y-axis represents the different models used, the x-axis represents the number of entries evaluated. Red indicates false negative, orange denotes false positive, light green signifies true negatives, and dark green indicates true positives. In this case, a true positive indicates a record the language model correctly labeled as a positive, a true negative indicates a record the model correctly labeled as a negative, a false positive is a record the model erroneously labeled as a positive, and a false negative a record that the model erroneously labeled as a negative. Bar heights and error bars indicate the mean and standard deviation of n = 3 replicates, respectively. In some cases, a model-prompt pair generated the same outcome in all three replicates, leading to standard deviation values of zero and thus appearing in this figure as a bar without an error bar.

With a collection of manually validated positive and negative records in hand, we next retrieved the full contents of each record, including the PubMed IDs of any articles describing the enzyme in the record. In a separate step, we retrieved the abstracts of the articles with those PubMed IDs and associated those abstract with the corresponding records in our collection. Thus, at this stage, our collection consisted of the 142 records, each with their Protein ID, full record contents, associated abstracts, and manual annotation as a positive or negative record (Fig. 1A). Finally, we looped over all the records in our collection and passed them 1-by-1 to an OpenAI language model alongside a system prompt designed to elicit a reponse from the model labelling the record as positive or negative. We compared the label assigned by the language model against our manually assigned label, and, considering the manually assigned label to be factual, labeled each language model response as a true positive, true negative, false positive, or false negative. In this case, a true positive indicates a record the langauge model correctly labeled as a positive, a true negative indicates a record the model correctly labeled as a negative, a false positive is a record the model erroneously labeled as a positive, and a false negative a record that the model erroneously labeled as a negative.

Using the approach above, we tested the accuracy of several language models under the instruction of several different styles of system prompts. A system prompt is a model input that describes a task that the language model should perform. System prompts can range from very simple to very complex and have a substantial impact on the efficacy of the model (22–24). Furthermore, through the process of prompt engineering, a prompt can be optimized to increase the system’s performance against a given metric (25). A key example emphasizing the importance of prompt engineering is the “chain-of-thought” prompting technique developed in 2022 (9). The most basic form of the method consists of simply adding the words “Lets think through this step-by-step” into an existing prompt. The researchers found that by adding these words to a prompt, model performance increased substantially (from 18% to 80% accuracy on the MultiArith test using OpenAI GPT3) (9). The chain-of-thought prompting method has since become a standard prompting technique for increasing language model performance, to the extent that OpenAI has incorporated it as a default behavior into some of its most recent models (26). Here, we evaluated the accuracy of four different language models (gpt-3.5-turbo, gpt-4o-2024-08-06, gpt-4-0125-preview, and gpt-4o-mini) using one of three different system prompts (a standard, non-chain-of-thought prompt; a “let’s think step-by-step” chain-of-thought prompt, and a more detailed chain-of-thought prompt that asked the model to include intermediate steps in its output). Details of all system prompts used in this section are provided in Supplemental File 1, but all prompts were geared toward asking the model “Does the enzyme described in the record make the product?”.

After completing the tests described above, we noticed a number of interesting patterns in the results. Several patterns pertained to differences in performance between models. We observed that GPT-3.5 Turbo generated significantly more false negatives than the other models, regardless of the type of prompting strategy used. We also found that, generally speaking, the GPT-4 models were largely comparable. This suggests that model size, which is generally thought of as being correlated with model sophistication, is connected to the ability of the model to distinguish between true positive results and false negative results.

Next, we noticed several patterns in the results connected to the prompts that were used. Above all, we observed that when using the most sophisticated chain-of-thought prompt, the models were able to achieve a significantly higher rate of true negative detection, which seemed to be connected to a lower false positive rate (Fig. 1B). In other words, when using a sophisticated system prompt, all the models tested here were more frequently able to recognize instances in which an entry contained an enzyme that had been experimentally validated to make the associated product, rather than just predicted to make the product.

Considered together, our results so far suggest several major points. First, a general chain-of-thought prompt like “let’s think step by step” performs significantly worse than an explicit chain-of-thought prompt detailing all the steps that a model should take in generating its answer. In particular, it seems that this more detailed form of a chain-of-thought prompt leads the model to recognize false positives and correctly classify them as true negatives. This more detailed chain-of-thought prompt, in combination with newer, larger models, led to the highest performance, approaching 90% accuracy with regard distinguishing between validated versus predicted enzyme-product associations.

### B. Automated Prompt Engineering For Compound-Species Relationship Detection

Recognizing the critical role of prompt engineering in our analysis of extracting protein-product relationships from NCBI, our next objective was to explore how systematic prompt engineering and related strategies could enhance task-specific accuracy. In a previous study, we used a collection of >2500 manually evaluated abstracts to test the ability of large language models to identify text passages describing the identification of specific chemical compounds in extracts of particular plant species (13). The best-performing language model in that study successfully processed these text passages with a low false positive rate—indicating it rarely misattributed a specific chemical to a plant species. However, this came at the expense of a high false negative rate, meaning the model often missed true instances of chemical production by particular species. Therefore, in this section, our goal was to determine if prompt engineering and related strategies could improve this high false negative rate.

For systematic prompt engineering, we implemented a method that leveraged language models to generate a diverse array of prompts. This method involved two language models: the first, termed the prompt generator language model, was responsible for generating prompts, while the second, the prompt evaluator language model, assessed the generated prompts against cases with established answers. Initially, the prompt generator language model created 50 prompts, which we then tested for accuracy using the evaluator language model and 20 test cases. The accuracy of these prompts ranged from 65% to 90%, with the majority achieving an accuracy of either 75% or 80% (Fig. 2A, inset histogram).

**Fig. 2.**
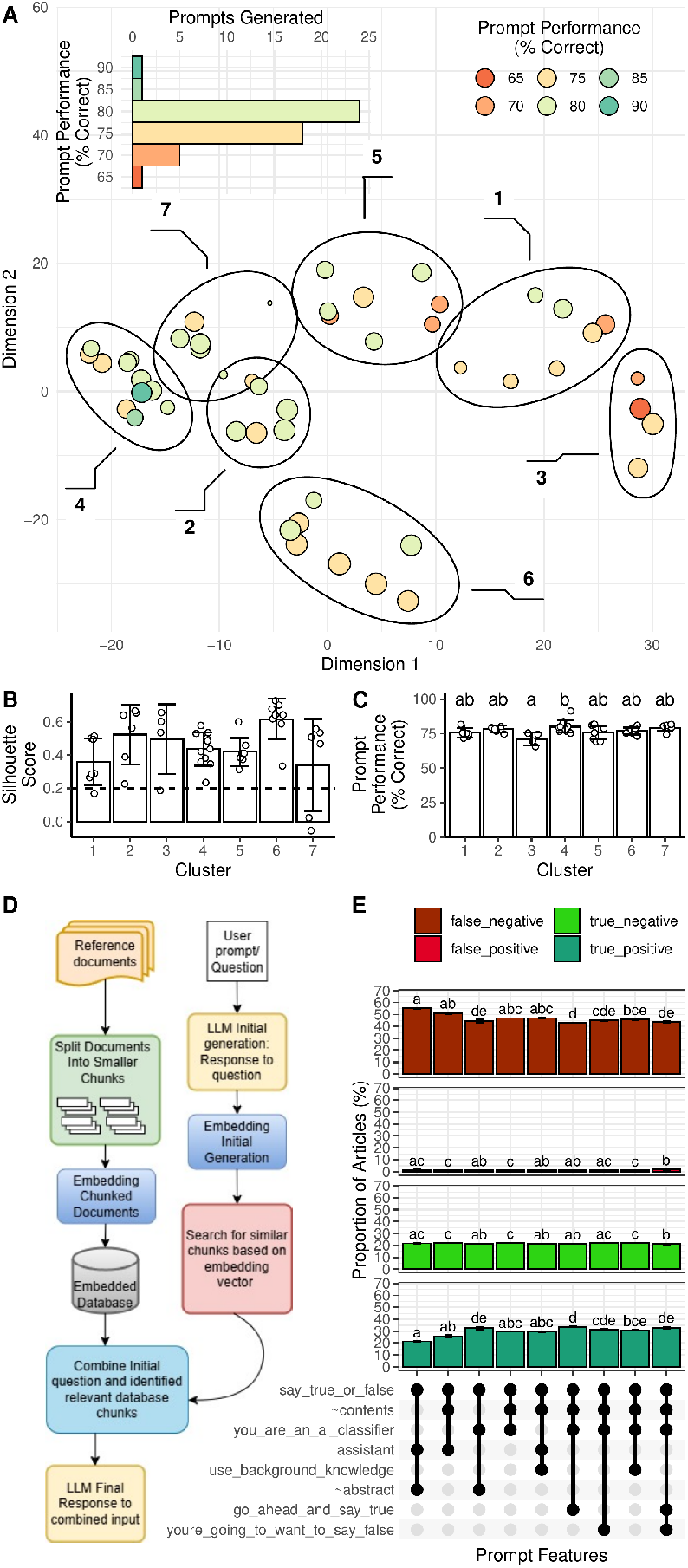
Automated prompt engineering and retrieval-augmented generation to enhance the identification of compound-species associations. **A** Performance (inset histogram) and principal components analysis of the linguistic meaning of 50 prompts generated for the task of compound-species relationship identification. Ellipses indicate clusters of linguistically similar prompts. **B** Silhouette scores for each point in each cluster, where scores > 0.5 suggest strong cluster membership, scores between 0.2 and 0.5 indicate moderate support, and scores below 0.2 show low cluster association. **C** Performance of the prompts in each cluster. Bar heights and error bars indicate means and standard deviations of n = 3 replicates, respectively. Letters indicate statistical groups determined by Tukey’s HSD test (adjusted p < 0.05). **D** Model of retrieval-augmented generation. **E** Performance of different system prompts in combination with retrieval-augmented generation. Contents of the system prompts are indicated on the x-axis. Colors indicate true/false positives/negatives, as described for Figure 1.

To investigate the reasons behind the observed differences in prompt performance, we employed an embedding model to generate numeric vectors representing the semantic content of each prompt. We then applied principal component analysis to these vectors, allowing us to map the prompts within a two-dimensional space defined by the first two principal components (Fig. 2A). Using k-means clustering, we identified clusters within this principal component space. Silhouette scores, which measure how similar each point is to others within its cluster compared to those in other clusters, indicated that a seven-cluster model provided a reasonable fit for the data distribution in the principal component space (Fig. 2B). Interestingly, this linguistic-based separation in principal component space also correlated with performance outcomes: high-performing prompts predominantly clustered on the left side, while lower-performing prompts clustered on the right. In particular, the leftmost cluster (Cluster 4) and the rightmost cluster (Cluster 3) demonstrated significantly different performance levels (Wilcoxon test, p < 0.05). These results suggest that linguistic variations in prompt phrasing can lead to substantial performance differences, even among prompts designed for the same task, a phenomenon that has been reported by others (12, 27). Together, our results and those in the literature strongly suggest that a diverse range of task-specific prompts should be tested in language model applications before deploying a large-scale pipeline.

Alongside the prompt engineering experiments described above, we also evaluated a well-known language model technique called retrieval-augmented generation (RAG) (28, 29). Retrieval-augmented generation enhances language model outputs by supplying domain-specific information alongside the input prompt. A basic implementation of RAG involves creating a system where a user poses a question, and the system retrieves relevant excerpts from a content database to provide as additional context to the language model for response generation (Fig. 2D). In our study, we used the manually evaluated abstracts from a prior study as the content database (“literature,” Fig. 2D, (13)) from which the RAG system retrieved information to use alongside each question as input to the language model. For example, we asked questions like “Is compound X produced by species Y?” and included the abstract excerpts retrieved by the RAG system as part of a structured prompt to the language model to assess the question. This approach not only provides the language model with clear instructions through a detailed system prompt but also supports response generation with peer-reviewed information from the content database. Full prompts used in this section are provided in Supplemental File 2.

Using the retrieval augmented generation system described above, along with a set of modular system prompts, we evaluated the performance of the language model GPT-4o. For each combination of content (abstract or retrieved content) and system prompt, we evaluated the model over 200 compound-species associations for which we had previously manually determined the answers. We then compared the model’s responses against the manually determined correct answers and calculated true positive, false positive, true negative, and false negative rates for each prompt and content combination. The largest effect we observed was when the system prompt provided a detailed description of the role that the model should play. For example, instructing the model that it was simply a “helpful assistant” yielded results similar to what we had observed previously, with a 55% false negative rate (13). However, when we told the model to act as an “AI classifier” and provided further details about how it should perform that role, we saw a significant reduction in the false negative rate (down to ∼40% from ∼55%). Others have reported on the importance of role- and goal-oriented prompts in creating effective language model pipelines (30). We did detect some smaller effects due to differences in providing abstract versus retrieved content or adding additional modular components in the system prompt that attempted to encourage the model to describe compound-species relationships as true (an attempt to reduce the false negative rate). However, these effects were minor compared to the impact of providing a well-defined role as part of the system prompt. Thus, taken together with the literature, our experiments suggest that providing additional, perhaps redundant, content to the model does not substantially improve the false negative rate. However, prompt engineering and prompt specifics have a substantial and significant impact. We suggest that future tests include styles of system prompt engineering that involve few-shot and multi-shot learning. These are prompting styles in which examples of how to handle common and edge cases are provided to the model, increasing its ability to accurately handle task-specific activities.

### C. Vision AI For Quantitative Data Table Transcription

Our final objective was to assess the potential of multimodal language models to generate structured data relevant to plant metabolic research. Substantial amounts of quantitative data on phytochemical profiles are stored in tables within scientific articles. In some instances, these tables are provided in a machine-readable format, such as CSV files in supplementary materials. However, such files are not always provided, particularly not by older publications. Consequently, tables embedded in PDF files represent a valuable yet underutilized source of information that is labor-intensive to systematically compile into a larger structured dataset for analysis. Without developing automated or semi-automated methods to unify this information, we risk losing the results gathered over decades of research.

To evaluate the ability of multimodal language models to transcribe quantitative data from tables published in scientific articles, we began by collecting 13 complex tables from a set of 13 different scientific articles describing the abundances of phytosterols in the seed oils of various plant species (32–44). The tables in these articles reported the abundances of phytosterols in 67 different plant species, with many species being described by multiple tables in the set. Initially, we attempted to use the multimodal model to extract data from PNG images of these table images directly, but we found that reproducible and accurate transcription was severely hindered by the non-uniform structure of the tables. In their raw form, many of the tables included units of scientific measurements in row names, reports of ranges rather than means and standard deviations, and complex layouts with summary statistics or multiple presentations of the same data in different units, including percentages. The poor performance of the model in this environment of non-uniformity led us to explore strategies to improve accuracy.

To improve table transcription by multimodal language models, we first manually simplified the images by importing the tables into image editing software and removing portions that appeared to confuse the model. These included table captions and rows or columns containing summary statistics. We also replaced the common names of plant species in each table with their latin names as reported in respective articles, then saved each table as a high-resolution PNG image. Next, we divided the transcription process into two phases (Fig. 3A). The first phase was image extraction, where PNG images of each table were encoded as base64 images and then passed to a multimodal language model (OpenAI gpt-4o) that transcribed the table as a CSV file. The resulting CSV data was then passed to a second language model to verify the presence of chemical names or plant species names. If no recognizable names were present, the multimodal model was prompted to attempt the transcription again. Otherwise, the CSV file containing both compound and species names was written to disk and, if necessary, automatically transposed so that compound names were organized as row headers and species names as column headers.

**Fig. 3.**
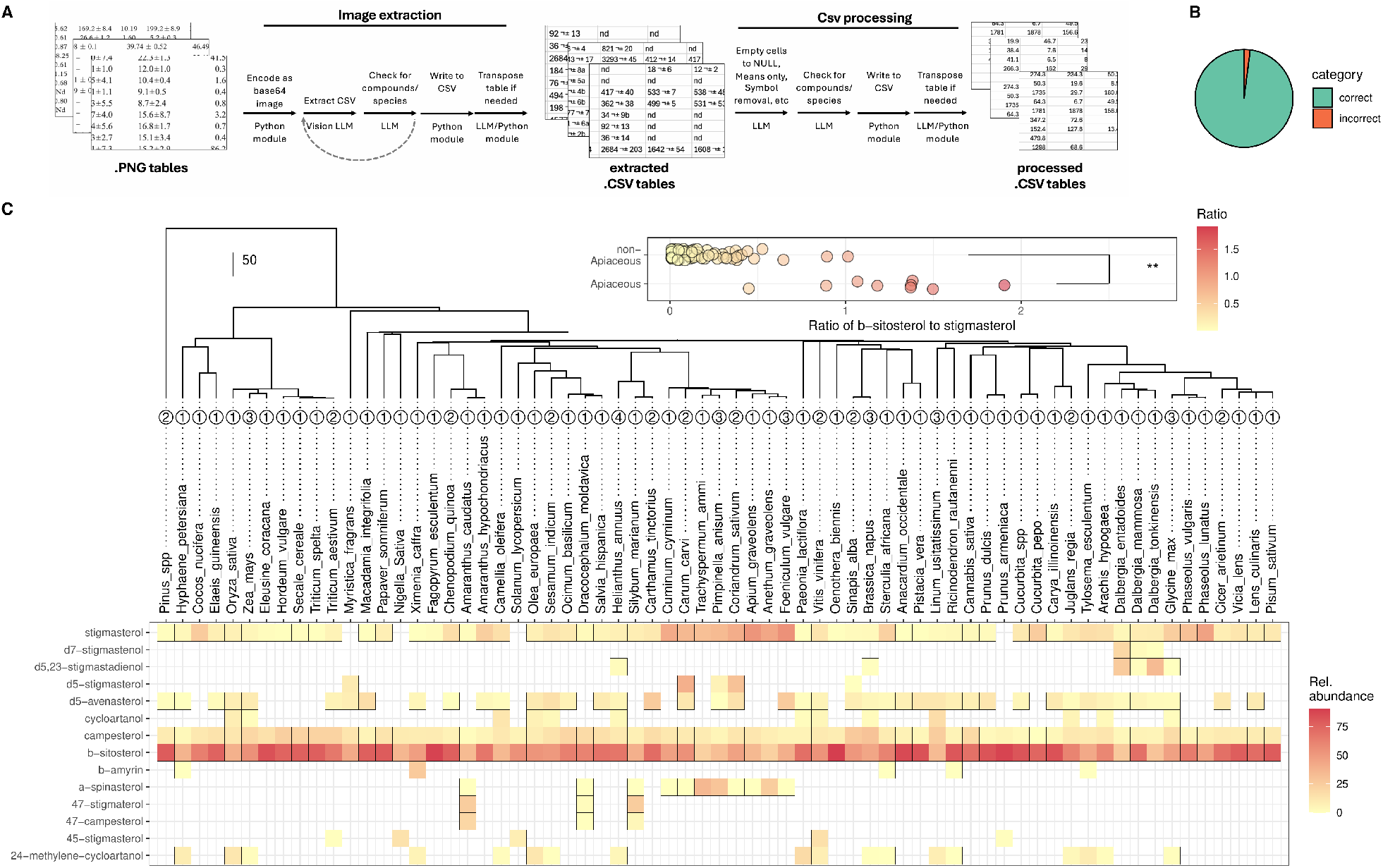
Workflow and results from using a multimodal language model to transcribe images of scientific tables. **A** Workflow used to transcribe/extract and processes text from images of tables. **B** Pie chart indicating the accuracy of the transcription process based on manual comparison of transcribed data and the original tables. **C** Phylochemical map showing the abundance (in heat map, abundance indicated in legend) of different phytosterols across the species included in this study (in phylogenetic tree). The phylogeny was prepared by pruning a previously published megaphylogeny (31). The inset plot shows the ratio of beta-sitosterol to stigmasterol in Apiaceous versus non-Apiaceous species. Stars indicate statistically significant differences in the mean ratio between the two groups (Wilconxon test, p < 0.05).

In the second phase of our pipeline, each CSV file was processed using a combination of language model calls and Python modules (Fig. 3A). First, each CSV file was read from the disk and passed to a language model that retranscribed the table in a simpler format that omitted symbols, standard deviations, and other non-standard characters. The output from that language model was then checked by another language model, which verified the presence of chemical compound names and species names before writing the table to a CSV file and conducting transposition if necessary. This sequence of steps resulted in processed CSV tables, which were then manually compared against the original images. This manual comparison revealed that the processed tables were 98% accurate compared to the original images (Fig. 3B). Overall, our pipeline and its results suggest that, though some manual intervention is required, multimodal models are highly useful for creating structured data from previously reported research results.

To illustrate one possible application of (semi-)automated table transcription by large language models, we next placed the transcribed quantitative data into a phylogenetic context. We pruned a megaphylogeny (31) to create a phylogeny for the 67 species reported across the transcribed tables. We then created a heat map next to the tree showing the relative abundance of different phytosterols in the seed oils of those species, based on the table data (Fig. 3C). This revealed that, in most species, beta-sitosterol was the most abundant phytosterol. However, in some species, stigmasterol was also a prominent component. We examined the ratio of stigmasterol to beta-sitosterol in each species for which both compounds had been reported. Interestingly, transcribed data from three different publications consistently showed that species in the carrot family (Apiaceae) had significantly higher ratios of stigmasterol to beta-sitosterol, suggesting that a lineage-specific event may have occurred in the ancestor of this group, leading to altered phytosterol ratios in seed oils (Fig. 3C, inset).

While developing the multimodal language model-based transcription pipeline, we made several observations that will likely impact future implementations of similar systems. Perhaps our most important observation was the considerable variation in table structure between publications. Tables varied greatly and often included forms of normalized or summarized data interspersed with abundance measurements. A primary challenge in developing this pipeline was determining how to systematically distinguish between abundance values and other types of information in table cells. In the end, we overcame this difficulty through manual annotation.

Having to manually inspect and polish the tables is perhaps beneficial because it forces human intervention and makes the researcher notice details like which solvents were used, whether different plant varieties were tested, or if various growing conditions were applied. Being aware of this sort of data for the species or chemicals of interest seems advantageous, albeit at the cost of throughput.

We also noticed that using high-resolution images greatly increased the accuracy of language model transcription. Low-resolution images were almost always transcribed incorrectly, with inaccuracies in both numerical transcription and the assignment of particular numbers to specific table cell positions. Interestingly, we found that table size did not seem to impact accuracy; processing a 3×5 table was just as accurate as transcribing a 10×14 table, provided the tables’ images are of sufficient resolution. This suggests that, as long as highresolution images can be obtained, this workflow should be accurate for large table sizes. However, scaling this pipeline to handle a large number of tables will require automated or semi-automated solutions for managing the non-uniformity of data types presented in quantitative tables.

## 4. Conclusions

Here, we conducted several explorations and evaluations of large language models in applications related to creating structured data from unstructured or largely unstructured information sources. Our key findings indicated that both prompt content at a general architectural level (standard prompts versus chain-of-thought prompts) and prompt content at a word level significantly affect the performance of a language model.

Carefully and specifically defining the role that a language model should play in the context of performing a particular task also has an impact on the model’s accuracy. When using a multimodal model, we noticed that high-resolution images led to higher transcription accuracy. However, complex table structures—including summary statistics and other information within a given table—led to difficulties in transcribing and merging multiple tables into a single unified format. Importantly, these difficulties, and our approach to overcoming them, highlight the importance of human involvement and subject matter expertise in pipelines that involve language models.

Overall, our results illustrate that large language models have considerable promise but are not without limitations. Several important observations include the task-specific manner in which prompts need to be designed for language models to perform with high accuracy and appropriately handle edge cases. Future implementations will need to be designed accordingly. We note that designing such prompts is a skill that will likely be useful in many disciplines and could serve as a valuable learning objective in relevant coursework (45). Similarly, collaboration between disciplines—including plant sciences, linguistics, data science, and bioinformatics—could lead to advances in this area and facilitate the integration of language models with existing databases and tools that are important in plant metabolic research.

Looking forward, we note the growing trend in the literature of integrating language models with other related entities and approaches, including machine learning and knowledge graphs (46). It is also interesting to note the reported efficacy of small (as opposed to large) language models in highly discipline-specific tasks (47–49). The development of multimodal models may also influence our field in the future as models capable of interpreting both text and biological sequence data are developed (50, 51). In all these cases, the importance of labeled data is clear, and we would like to highlight the need to support efforts geared toward generating such data. Creating such datasets can help us not only use existing language models most effectively in our discipline but could also pave the way for the creation of discipline-specific models that perform at high levels and support the advancement of plant metabolic research.

## 5. Supplemental Materials

Supplemental File 1: Prompts used in identifying enzyme-product pairs. Supplemental File 2: Prompts used in identifying compound-species pairs. All other code, prompts, and raw data are available in the project repository at https://github.com/thebustalab/ai_in_phytochemistry.

## 6. Acknowledgements

The authors wish to acknowledge the support of Alan Oyler during this project. We also wish to acknowledge the University of Minnesota Duluth Chemistry and Biochemistry Department, the UMD Undergraduate Research Opportunities Program, the UMD Summer Undergraduate Research Program, and the UMD Chemistry Master’s program for providing the funding and facilities to make this research possible. During the writing of this manuscript, the large language model GPT-4o was utilized for copy editing to suggest alternative sentence structures and word choices that enhanced grammar and readability. Finally, we collectively acknowledge that the University of Minnesota Duluth is located on the traditional, ancestral, and contemporary lands of Indigenous people. The University resides on land that was cared for and called home by the Ojibwe people, before them the Dakota and Northern Cheyenne people, and other Native peoples from time immemorial. Ceded by the Ojibwe in an 1854 treaty, this land holds great historical, spiritual, and personal significance for its original stewards, the Native nations, and peoples of this region. We recognize and continually support and advocate for the sovereignty of the Native nations in this territory and beyond. By offering this land acknowledgment, we affirm tribal sovereignty and will work to hold the University of Minnesota Duluth accountable to American Indian peoples and nations.

## 7. Data Availability Statement

All data and code used in this study are available for free download from the project repository at: https://github.com/thebustalab/ai_in_phytochemistry. The Supplemental Files contain all the prompts used in identifying enzyme-product and compound-species pairs.

