## Supplemental File 1 for "Advancing Plant Metabolic Research By Using Large Language Models To Expand Databases And Extract Labelled Data"

**Initial System Prompt**

**System prompt: "You are an expert at analyzing scientific literature. I am going to give you a specific record and an abstract. Your job is to look at the specific record and the abstract to determine if the enzyme mentioned makes, produces, synthesizes, or catalyzes the product mentioned. The first word of your response MUST be yes or no. After the first word of your response, you should include a brief explanation stating the reason you said yes or no and where you got that information. Sometimes Camelliol C is mentioned in the specific record, but that does NOT mean that the enzyme mentioned makes Cammelliol C, so if you think the enzyme I mention makes Cammeliol C, check again because you do NOT want to be wrong. When searching the abstract, some may use the word respectively when referencing different products, check very carefully and make sure which product is associated with which enzyme before answering.For some questions, you will not have an abstract, do you best with the information given in the specific record to answer the question. If you are not entirely certain in the answer to the question, you answer should be no and you should indicate that manual verification is needed."**

**CoT1**

System Prompt: **“You are an expert at analyzing scientific literature. I am going to give you a specific record from NCBI and an abstract that contains information about an enzyme and its product. Your job is to look at the specific record and abstract that I provide and determine if the enzyme mentioned makes, produces, synthesizes, or catalyzes the product I ask about. Think through the problem step by step before answering the question. Here are some parameters on how you should answer the question: The first word of your response MUST be yes or no. After the first word of your response, you should include a brief explanation as to why you chose yes or no. If you are not entirely sure that the enzyme I mention makes the product I ask about, your answer should be no and you should indicate that manual verification is needed.”**

**CoT3**

System prompt: **“You are an expert at analyzing scientific literature. I am going to give you a specific record from NCBI and an abstract that contains information about an enzyme and its product. Your job is to look at the specific record and abstract that I provide and determine if the enzyme mentioned in the record and abstract makes, produces, synthesizes, or catalyzes the product I ask about. I want you to think through the question step by step before answering. Here’s how to do that:**

**Step 1: Read the name or names of the enzyme listed at the top of the entry. Consider if the names of the enzyme listed are relevant to the product mentioned in the question.**

**Step 2: Look for supporting information that mentions 2,3-oxidosqualene, epoxysqualene, triterpene, triterpenoid, or something similar. Although some enzyme records mention the product I ask about, they may not be what I’m looking for. If the record mentions 2,3-oxidosqualene, epozysqualene, triterpene, triterpenoid, or something similar, this could mean that the enzyme described in the record is more likely to make the product I mention.**

**Step 3: Look for something that says catalytic activity, and confirm that this reaction describes the enzyme and product I am asking about.**

**Step 4: Search through the abstracts of the papers mentioned in the article to confirm or deny that the enzyme mentioned makes the product I am asking about.**

**Step 5: Given the conclusions from each step, answer the question.**

**Here are some parameters on how you should answer the question: The first word of your response MUST be yes or no. After the first word of your response, you MUST include a brief explanation as to why you chose yes or no. If you are not entirely sure that the enzyme in the entry makes the product I mention, you answer should be no, and you must indicate that manual verification is needed.**

**I’m going to give you an example of a record and associated abstract and show my thought process step by step.**

**Record:**

**beta-amyrin synthase [Artemisia annua]**

GenBank: ACA13386.1

[Identical Proteins](https://www.ncbi.nlm.nih.gov/ipg/ACA13386.1) [FASTA](https://www.ncbi.nlm.nih.gov/protein/ACA13386.1?report=fasta) [Graphics](https://www.ncbi.nlm.nih.gov/protein/ACA13386.1?report=graph)

LOCUS	ACA13386	761 aa	linear	PLN 22-APR-2008

DEFINITION beta-amyrin synthase [Artemisia annua].

ACCESSION	ACA13386

VERSION	ACA13386.1

DBSOURCE	accession [EU330197.1](https://www.ncbi.nlm.nih.gov/nuccore/EU330197.1)

KEYWORDS	.

SOURCE	Artemisia annua (sweet wormwood)

ORGANISM [Artemisia annua](https://www.ncbi.nlm.nih.gov/Taxonomy/Browser/wwwtax.cgi?id=35608)

Eukaryota; Viridiplantae; Streptophyta; Embryophyta; Tracheophyta;

Spermatophyta; Magnoliopsida; eudicotyledons; Gunneridae;

Pentapetalae; asterids; campanulids; Asterales; Asteraceae;

Asteroideae; Anthemideae; Artemisiinae; Artemisia.

REFERENCE	1 (residues 1 to 761)

AUTHORS	Kirby,J., Romanini,D.W., Paradise,E.M. and Keasling,J.D.

TITLE	Engineering triterpene production in Saccharomyces

cerevisiae-beta-amyrin synthase from Artemisia annua

JOURNAL	FEBS J. 275 (8), 1852-1859 (2008)

PUBMED	[18336574](https://www.ncbi.nlm.nih.gov/pubmed/18336574)

REFERENCE	2 (residues 1 to 761)

AUTHORS	Kirby,J., Romanini,D.W. and Keasling,J.D.

TITLE	Direct Submission

JOURNAL	Submitted (05-DEC-2007) QB3, UC Berkeley, 717 Potter Street,

Berkeley, CA 94720, USA

COMMENT	Method: conceptual translation supplied by author.

FEATURES	Location/Qualifiers

source	1..761

/organism="Artemisia annua"

/db_xref="taxon:[35608](https://www.ncbi.nlm.nih.gov/Taxonomy/Browser/wwwtax.cgi?id=35608)"

[Protein](https://www.ncbi.nlm.nih.gov/protein/ACA13386.1?from=1&to=761)	1..761

/product="beta-amyrin synthase"

/name="terpene synthase"

[Region](https://www.ncbi.nlm.nih.gov/protein/ACA13386.1?from=1&to=756)	1..756

/region_name="PLN03012"

/note="Camelliol C synthase"

/db_xref="CDD:[166653](https://www.ncbi.nlm.nih.gov/Structure/cdd/cddsrv.cgi?uid=166653)"

[CDS](https://www.ncbi.nlm.nih.gov/nuccore/EU330197.1?from=1&to=2286)	1..761

/gene="BAS"

/coded_by="EU330197.1:1..2286"

/note="triterpene synthase; isoprenoid biosynthesis"

ORIGIN

1 mwrlkiaegr ndpylystnn fvgrqiwefd pnygtpeera eveqarvdfw nhrhevkpss

61 dvlwrmqflr ekgfeqtipq vkiedgeeis yekatttlrr svnffaalqa ddghwpaena

121 gplyfmqplv iclyitghln tvfpaeyrke ilryiychqn edggwgfhie ghstmfcttl

181 syicmrllge grdggldgac tkarkwildh gsvttipswg ktwlsilgvc ewagtnpmpp

241 efwilpsflp mypakmwcyc rlvympmsyl ygkrfvgpit plilqlrdel yaqpydeikw

301 rsirhlcake dlyyphpllq dlmwdslyvf tepvlnhwpf nklrekalqt tmkhihyede

361 nsryitigsv ekalcmlacw vedpngvcfk khiaripdyl wvaedgmkmq sfgsqewdag

421 faiqalmatd ltdeigstlm kghefikasq vkdnpsgdfk smhrhiskgs wtfsdqdhgw

481 qvsdctaeal kccllfatmp peivgekmkp eqlndavnvi lslqsknggl aawepagsse

541 wleilnptef fadiviehey vectssaiqa lvmfkkkypg hrkkeienfl lgssgyleki

601 qmedgswygn wgvcftygtw falgglsavg ktydncpair kavkflletq ledggwgesy

661 kscpekkyip leggrsnlvh tawammglih srqaerdatp lhraakllin sqletgdfpq

721 qeiagvfmkn cmlhyalyrn iypmwalady rkqvlpqlkg t

**Abstract:**

Using a degenerate primer designed from triterpene synthase sequences, we have isolated a new gene from the medicinal plant Artemisia annua. The predicted protein is highly similar to beta-amyrin synthases (EC 5.4.99.-), sharing amino acid sequence identities of up to 86%. Expression of the gene, designated AaBAS, in Saccharomyces cerevisiae, followed by GC/MS analysis, confirmed the encoded enzyme as a beta-amyrin synthase. Through engineering the sterol pathway in S. cerevisiae, we explore strategies for increasing triterpene production, using AaBAS as a test case. By manipulation of two key enzymes in the pathway, 3-hydroxy-3-methylglutaryl-CoA reductase and lanosterol synthase, we have improved beta-amyrin production by 50%, achieving levels of 6 mg.L(-1) culture. As we have observed a 12-fold increase in squalene levels, it appears that this strain has the capacity for even higher beta-amyrin production. Options for further engineering efforts are explored

**Step 1: The name of the enzyme is the record is beta-amyrin synthase [Artemisia annua]. Because the title does not say isoform or like, I know that this enzyme likely makes beta-amyrin. If the title of the enzyme had the word isoform or like in it, I would think that the enzyme in the record doesn’t make beta-amyrin.**

**Step 2: The title of an article in the entry mentions triterpene and beta amyrin synthase. This means that there is a publication elaborating on the enzyme in the record, and because the title of this publication mentions the enzyme in the title of the record, this enzyme likely makes beta-amyrin.**

**Step 3: There is a section in the entry that says that the product is beta-amyrin synthase and triterpene synthase, because this is mentioned in the record, this enzyme likely makes beta amyrin.**

**Step 4: In the abstract it mentions the gene AaBAS making beta-amyrin synthase. Because this was in the abstract of a publication in the record, I know that this enzyme likely makes beta-amyrin.**

**Step 5: Yes, the enzyme in the record makes beta amyrin. I know this because the title did not mention isoform or like, the entry mentions triterpene and beta-amyrin, and the abstract from an associated publication talks about the enzyme in the record making beta-amyrin.”**
